# The effect of motor-induced shaft dynamics on microtubule stability and length

**DOI:** 10.1101/2022.05.04.490659

**Authors:** Joël Schaer, Mireia Andreu-Carbó, Karsten Kruse, Charlotte Aumeier

## Abstract

Control of microtubule abundance, stability, and length is crucial to regulate intracellular transport as well as cell polarity and division. How microtubule stability depends on tubulin addition or removal at the dynamic ends is well studied. However, microtubule rescue, the event when a microtubule switches from shrinking to growing, occurs at tubulin exchange sites along the shaft. Molecular motors have recently been shown to promote such exchanges. Using a stochastic theoretical description, we study how microtubule stability and length depends on motor-induced tubulin exchange and thus rescue. Our theoretical description matches our *in vitro* experiments on microtubule dynamics in presence of kinesin-1 molecular motors. Although the average microtubule dynamics can be captured by an effective rescue rate, the dynamics of individual microtubules differs dramatically when rescue occurs only at exchange sites. Furthermore, we study in detail a transition from bounded to unbounded microtubule growth. Our results provide novel insights into how molecular motors imprint information of microtubule stability on the microtubule network.

**SIGNIFICANCE:** The microtubule network is essential for vital cellular processes like the organization of intracellular transport and division. Although microtubule assembly occurs at its tips, it has recently been reported that tubulin is exchanged along the microtubule shaft. Tubulin exchange plays an essential role in regulating microtubule dynamics and can be induced by molecular motors. Here, we provide the first systematic study of the impact of shaft dynamics on the regulation of rescue events, where a microtubule switches from shrinking to growing. Our results illustrate how the usage of microtubules as tracks for intracellular transport regulates the microtubule network and thus offers a novel perspective on intracellular organization.

## INTRODUCTION

In cells, the microtubule network underlies intracellular transport, segregates chromosomes during division, and is involved in establishing as well as maintaining cell polarity (1). To accomplish these diverse biological functions, dynamic properties of single microtubules within the network are tightly regulated determining their stability, length and local abundance (2). Microtubules grow by addition of tubulin bound to Guanosine-triphosphate (GTP) at their ends. The GTP is subsequently hydrolyzed to GDP, which affects tubulin affinity for microtubule binding and drives microtubule dynamics out of thermodynamic equilibrium (3). With GDP-tubulin at the tip, microtubules stop growing and start to shrink by tubulin removal from the tip, an event called catastrophe (4). Conversely, microtubules can stop shrinking and re-grow in a so-called rescue event (4).

Growth, shrinkage, and catastrophes can be regulated at the plus end. For example, the microtubule polymerase XMAP125 accelerates growth (5–7). This can also be achieved by the Cytoplasmic Linker Protein of 170 kDa (CLIP-170), which simultaneously enriches GTP-tubulin at the growing tip by binding to end-binding proteins 1 and 3 (EB1 and EB3) (8). Furthermore, interaction of EB1 and EB3 with the growing microtubule tip increases the catastrophe rate (9, 10). As a last example, the rate of tubulin removal at the tip can be affected by depolymerizing molecular motors like kinesin-8 (11), kinesin-13 (MCAK) (12), or kinesin-14 (Kar3p) (13). Since the abundance of molecular motors on a microtubule increases towards the tip, the regulation of tubulin removal by molecular motors can lead to an effectively length-dependent disassembly rate (14). Similarly, modifications along the microtubule shaft have been shown to induce a length-dependent depolymerization rate (15–17). Theoretical studies have analyzed in detail the implications of a length-dependent depolymerization rate on the microtubule length distribution (18–23). Let us also note that in addition to microtubule length regulation at the tip, severing provides a versatile mechanism to regulate microtubule length (24, 25).

The impact of rescue on microtubule dynamics is much less studied, because the underlying mechanism was unknown for a long time. Only recently it has been discovered that tubulin can exchange all along the microtubule shaft (26) leading to ‘GTP-tubulin islands’ (27). The molecular mechanism responsible for island formation is not yet understood (28), but mechanical forces and severing enzymes like spastin and katanin can enhance tubulin exchange (26, 29–31). Microtubules stop shrinking and resume growth preferentially at such islands (29, 30, 32, 33). It was found that the molecular motor kinesin-1 can damage the microtubule shaft and thus induce incorporation of GTP-tubulin (34–36). Through this process, kinesin-1 motors walking along microtubules increase the probability of rescue events and thus microtubule stability (35).

In the present work, we study theoretically how molecular motors increase microtubule length by regulating rescue events. To this end, we use stochastic simulations and theory and compare our results to *in vitro* experiments. We find that beyond a critical motor concentration, microtubule growth is unbounded (37, 38). Furthermore, we show that although effective rescue rates (37) capture well the average microtubule length in presence of molecular motors, the dynamics of individual microtubules changes significantly if tubulin exchange along the shaft is considered. In particular, subsequent growth and shrinkage phases are highly correlated in the entire regime of bounded growth. By including the effects of molecular motors on the shaft our model allows to study novel mechanisms of microtubule length regulation.

## MATERIALS AND METHODS

### Theoretical description of microtubule assembly kinetics

We represent a microtubule as a polar dynamic linear lattice, where the sites represent tubulin dimers, see Fig. 1. Each site is in one of three possible states, respectively denoted as T, D, and Tx, which correspond to GTP- and GDP-bound tubulin and to GTP-bound tubulin after exchanging GDP for GTP. A T-state switches to D at rate *k*_hyd_ and D turns into Tx at rate *k*_×_. The first of these transitions reflects spontaneous hydrolysis of a GTP bound to tubulin. The second transition accounts for the complex process of spontaneously replacing a GDP-tubulin from the microtubule lattice by incorporation of a GTP-tubulin. In our simulations, we refrain from describing this process in molecular detail. This is appropriate, because, in our experiments, we cannot resolve tubulin exchange on molecular time and length scales. Finally, Tx switches to D at a rate *k*_hydX_, which is in general different from *k*_hyd_.

**Figure 1:**
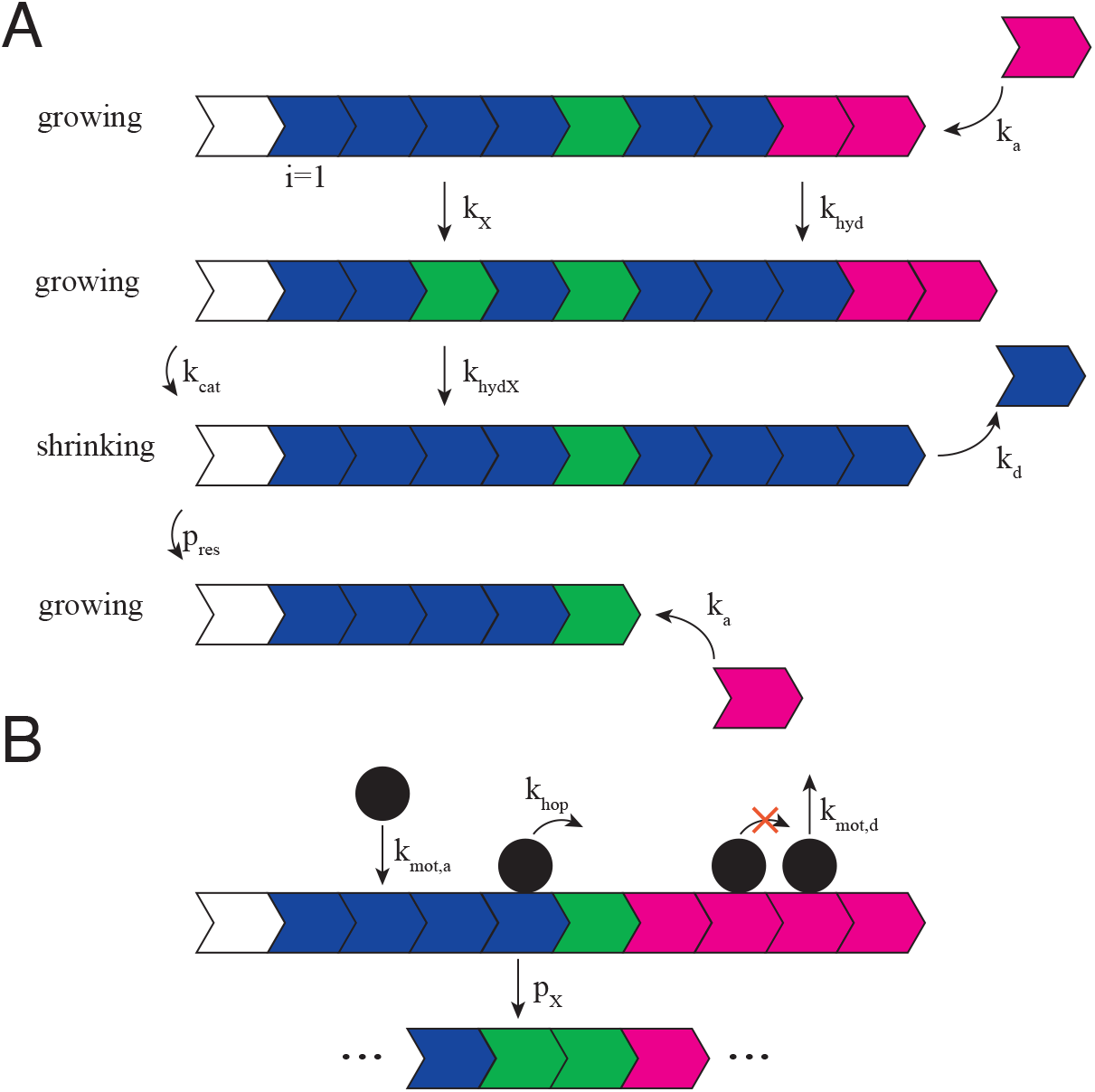
Schematic illustrations of theory and experiment. A) Microtubules are represented as linear polar lattices. Sites are in 3 different states: T (magenta), D (blue), Tx (green) corresponding to tubulin bound to GTP, GDP or to GTP after an exchange of GDP for GTP. Sites change form state T to D at rate *k*_hyd_, from D to Tx at rate *k*_×_, and from Tx to D at rate *k*_hydX_. The white site represents the inert seed. The lattice can be in a growing state, in which subunits in the T-state are added at rate *k*_*a*_, or in a shrinking state, in which subunits are removed at rate *k*_*d*_. The lattice switches from the growing to the shrinking state at rate *k*_cat_ and rescues with probability *p*_res_ if the site at the tip is in state Tx. B) Molecular motors are represented as particles. They attach to a free site at rate *k*_mot,a_ and detach from a site at rate *k*_mot,d_. Particles hop to an empty neighboring site towards the dynamic lattice end at rate *k*_hop_. If the particle leaves a D site, this site turns to the Tx-state with a probability *p*_×_.

The lattice as a whole can be in two dynamics states, namely, growing and shrinking. In the growing state, GTP sites are added at rate *k*_*a*_ at the plus end. In contrast, the minus end, is dynamically inert. This property reflects the use of stabilized microtubule seeds in our *in vitro* experiments. In the shrinking state, sites are removed from the plus end at rate *k*_*d*_ one by one. The transition of the lattice from the growing to the shrinking state corresponds to a catastrophe and occurs at rate *k*_cat_. In our *in vitro* experiments, we do not have access to the state of individual tubulin dimers. Consequently, in our description we take catastrophes to occur independently of the state of the site at the plus end. This is different from other theoretical studies, in which catastrophes are tightly linked to the loss of GTP-tubulin at the tip (39). Transitions from a shrinking to a growing lattice correspond to a microtubule rescue event. This event occurs with a probability *p*_res_ if the new site at the plus end is in state Tx, after removal of the preceding site.

We represent kinesin-1 molecular motors as particles hopping directionally on the lattice: at rate *k*_hop_, a particle moves to the neighboring site towards the plus end if this neighboring site is empty. This exclusion property accounts for steric interactions between molecular motors. We assume that detached motors form a reservoir such that particles attach to empty sites on the lattice at a constant rate *k*_mot,a_. In our experiments, this rate can be changed by changing the concentration of kinesin-1 in the buffer. In the simulations, we use 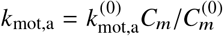, where *C*_*m*_ is the motor concentration and 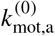 the motor attachment rate at 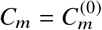. We take 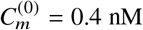, which was the smallest kinesin-1 concentration used in our experiments. Particles detach from the lattice at rate *k*_mot,d_.

Motor-induced exchange of GDP-tubulin for GTP-tubulin is captured by a probability *p*_×_ at which a site in state D is switched to Tx as a motor particle leaves the site by hopping. Similarly to the spontaneous exchange of a GDP-tubulin for a GTP-tubulin, this is an effective description of the potentially much more involved molecular process underlying motor-induced exchange of a GDP-tubulin for a GTP-tubulin.

### Stochastic simulations

Simulations of the model with **R**escue at **E**xchange **S**ites (RES-model) introduced above were carried out using a variant of the Gillespie algorithm, where initially the total rate *k*_tot_ of all possible processes is determined. After the time to the following event has been drawn from an exponential distribution with the characteristic time 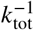, the identity of this event is determined according to the relative contribution of all possible events to the total rate. The simulations were performed using a custom-made c++ program.

Unless stated otherwise, the parameter values used in the simulations are listed in Table 1. We inferred the parameters directly from our *in vitro* experiments (35) whenever possible. To convert the observed microtubule growth velocity *v*_*g*_ into an attachment rate of the sites, we used the length of a single tubulin dimer, i.e., *d* = 8 nm, such that *k*_*a*_ = *v*_*g*_/*d*. In line with the literature, we chose *k*_hyd_ = 0.125 *s*^−1^ (40). Small changes of *k*_hyd_ didn’t substantially change the simulation results. We were left with 3 free parameters, *k*_hydX_, *p*_res_, and *p*_×_ that could not be obtained directly from our experiments or the literature. We determined their values by fitting the values of the rescue rate at 0 nM and at 0.4 nM kinesin-1 concentration to our experimental results.

**Table 1:** Parameter values used in the simulations.

| Parameter | Value |
| --- | --- |
| $k_a$ | $2 \text{ s}^{-1}$ |
| $k_d$ | $27 \text{ s}^{-1}$ |
| $k_{\text{cat}}$ | $0.005 \text{ s}^{-1}$ |
| $p_{\text{res}}$ | 0.15 |
| $k_{\text{hyd}}$ | $0.125 \text{ s}^{-1}$ |
| $k_{\text{hydX}}$ | $0.125 \cdot 10^{-3} \text{ s}^{-1}$ |
| $k_{\times}$ | $1 \cdot 10^{-5} \text{ s}^{-1}$ |
| $k_{\text{mot,a,0}}$ | $3.3 \cdot 10^{-5} \text{ s}^{-1}$ |
| $k_{\text{mot,d}}$ | $0.09 \text{ s}^{-1}$ |
| $k_{\text{hop}}$ | $16 \text{ s}^{-1}$ |
| $p_{\times}$ | 0.0018 |

### Meanfield equations

We introduce the mean values *t*_*i*_, *d*_*i*_, and *t*_×,*i*_ for site *i* = 1, 2, … being in state T, D, or Tx, respectively. These values range between 0 and 1 and we will refer to them as the probabilities of site *i* being in the respective states. Note that *F*_*i*_ ≡ [*t*_*i*_ + *d*_*i*_ + *t*_×,*i*_] is the probability of having a lattice with a length of at least *i* sites. We also introduce the mean values *g*_*i*_ and *s*_*i*_ for the growing and the shrinking lattice tips, respectively, to be at position *i* = 1, 2, … Finally, the mean value of having a motor particle at site *i* is denoted by *m*_*i*_.

Using a meanfield approximation, where correlations are expressed in terms of the mean values, the dynamic equations for the probabilities of the tip being at position *i* are

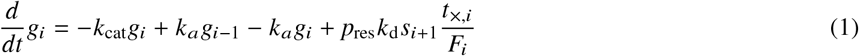

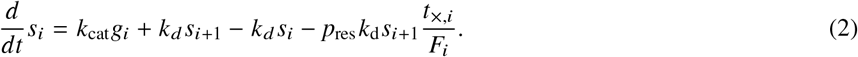

The first terms on the right hand sides account for catastrophes, the second and third terms for transitions due to addition or removal of a site. The final terms capture rescues, where *t*_×,*i*_/*F*_*i*_ is the fraction of sites *i* that are in the Tx state.

The dynamic equation for the probability being in state T is

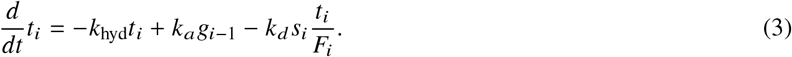

The term proportional to *k*_hyd_ represents the loss of a T-state by hydrolysis. The second term captures the addition of a T-state by polymerization at the growing tip, whereas the last term describes the loss of a T-state by site removal from the shrinking tip. Analogously, we have for the probability being in state D

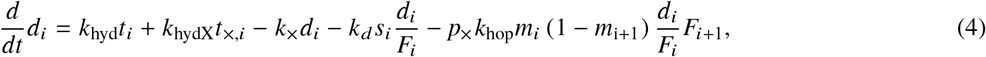

where the terms have an analogous meaning to those in Eq. (3). In addition, the terms proportional to *k*_hop_ account for transitions due to motor hopping. The dynamic equations for the probability being in state Tx follow the same logic. Explicitly, we have

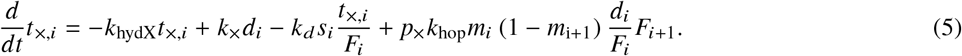

The dynamic equations are complemented by the boundary conditions

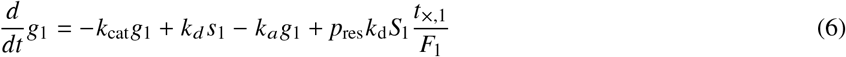

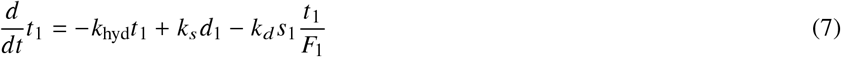

for Eqs. (1) and (3), whereas Eqs. (2), (4) and (5) hold also for *i* = 1.

Finally, we give the equation governing the evolution of the probability to find a motor at site *i* = 2, 3, …:

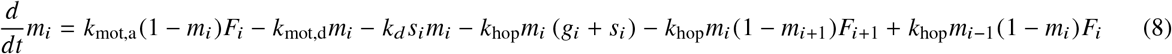

In this expression, the terms proportional to *k*_mot,a_ and *k*_mot,d_, respectively, account for motor attachment to and detachment from the lattice. The terms proportional to *k*_hop_ describe motor hopping towards the plus-end onto the neighboring site or off the lattice if at the tip. Finally, the term proportional to *k*_*d*_ captures motor loss due to depolymerization at the shrinking tip. The evolution of *m*_1_ is obtained from the above by setting *m*_0_ = 0.

### Distributions close to the seed

In case the lattice is very long, it is possible to simplify Eqs. (3)-(8) close to the seed by neglecting the tip dynamics. Explicitly, for sufficiently small values of *i*, we set *g*_*i*_ = 0 and *s*_*i*_ = 0, which implies *F*_*i*_ = 1. Furthermore, the probability to be in state T close to the seed is negligibly small and we set *t*_*i*_ = 0. In the continuum limit we then have

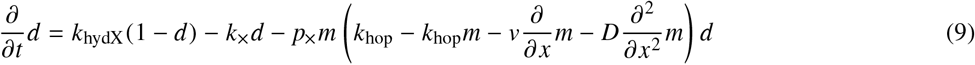

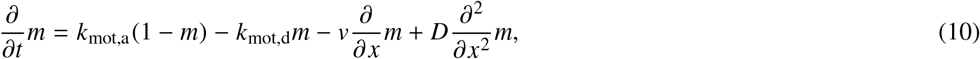

where we have used *t*_×_ = 1 − *d*. Furthermore, *v* = *ak*_hop_ and *D* = *a*^2^*k*_hop_/2 with *a* = 8.2 nm being the length of a lattice site corresponding to the length of a tubulin dimer.

### Microtubule assembly assay

Microtubules were polymerized using purified and labeled brain tubulin. Microtubules were polymerized in an assembled flow chamber and visualized under a TIRF microscope with a temperature stage controller stable at 37°C. A step-by-step protocol can be found in Ref. (35).

For microtubule dynamic assays and kymograph representations, microtubules were polymerized inside the flow chamber for 10 min at 37°C with 8 µM tubulin (20 % ATTO-565-labeled) in BRB80 1x containing anti-bleaching buffer (0.3 mg/ml glucose, 0.1 mg/ml glucose oxidase, 0.02 mg/ml catalase, 10 mM DTT, 0.125 % methyl cellulose, 1 mM GTP), 0.2 % BSA and 41.6 mM KCl. We injected to the polymerized microtubules either 0, 1 or 2 nM K430-GFP (kinesin-1) in presence of 8 µM tubulin (20 % ATTO-565-labeled) in BRB80 1x supplemented with anti-bleaching buffer, the chamber was sealed for imaging. Images were recorded every 3 to 4 seconds with the lowest laser power that allows microtubule visualization with reduced phototoxicity. To measure microtubule dynamics, kymographs were built with MultipleKymograph in ImageJ. We only analyzed isolated microtubules that did not bundle or cross with other microtubules.

For microtubule length studies, 8 µM tubulin (20 % labeled) and 0, 1 or 2 nM K430-GFP in BRB80 1x containing anti-bleaching buffer, 0.2 % BSA and 41.6 mM KCl were perfused inside the flow chamber and directly imaged at the microscope. Images from different positions were taken every minute for 45 min. We imaged only the microtubule channel to prevent as much as possible phototoxicity. Microtubule length was manually measured using ImageJ.

## RESULTS AND DISCUSSION

### Microtubule length distribution *in vitro*

We reconstituted *in vitro* microtubule growth in absence and in presence of the molecular motor kinesin-1. We limited our experiments to 45 min, because protein degradation started to affect microtubule dynamics after 45 min, and only considered isolated microtubules. In absence of kinesin-1, microtubules grew from stabilized seeds and showed dynamic instability with constant growth and shrinkage velocities, Fig. 2A. Once a microtubule started shrinking, we rarely observed rescue events, such that a microtubule typically disassembled completely to the stabilized seed. Note that we do not consider regrowth from the seed to be a rescue event, but to be the nucleation of a new microtubule. The average microtubule length increased linearly, then slowed down and eventually saturated, Fig. S1A. In steady state, the average microtubule length is 7.3 ± 5.1 µm (mean ± std. dev), Figs. 2D, S1A,D.

**Figure 2:**
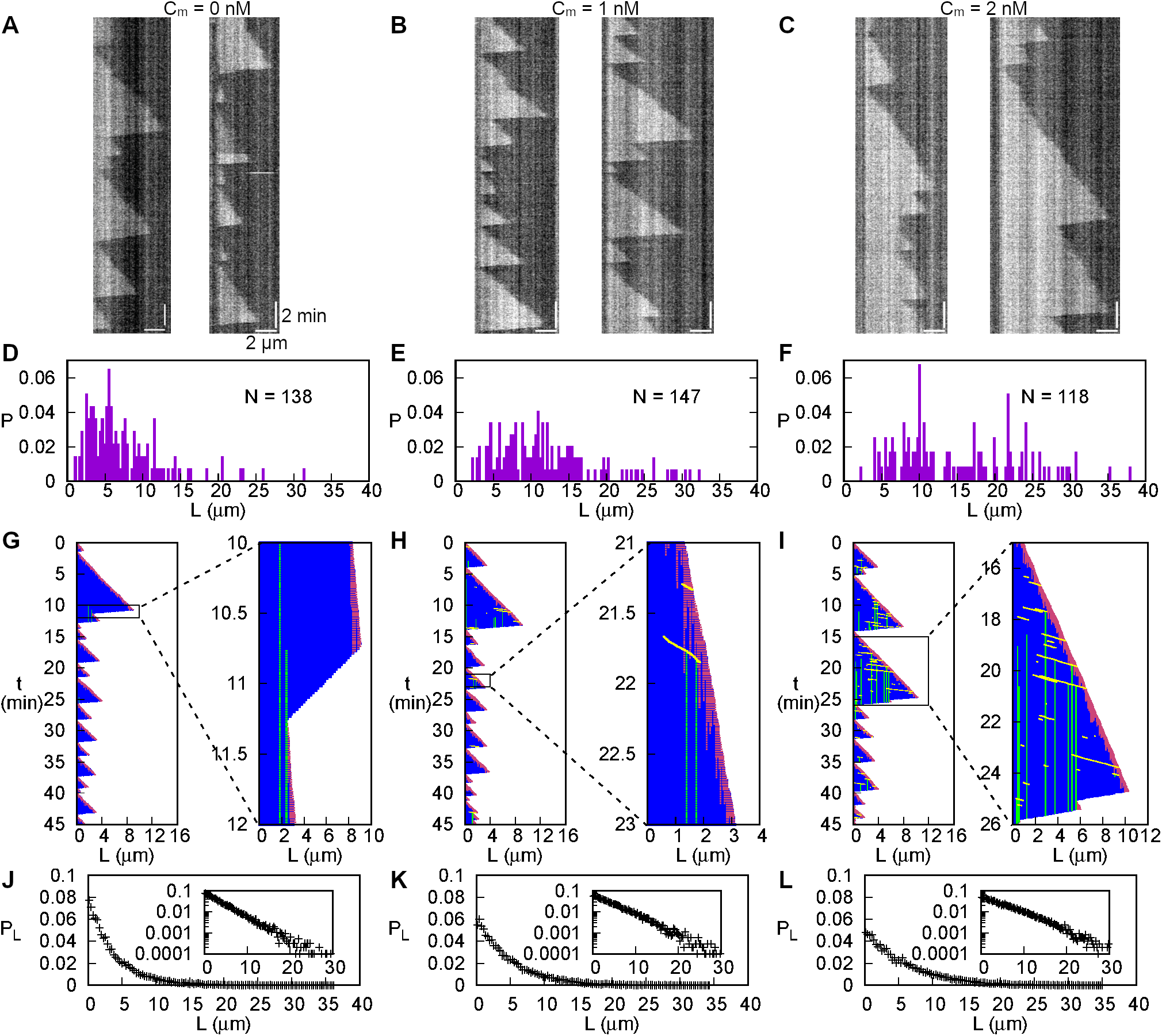
Measured and simulated microtubule dynamics in the presence of motors. A-C) Experimental kymographs for 0 (A), 1 (B), and 2 nM kinesin-1 (C). D-F) Experimental lengths distributions for 0 (D), 1 (E), and 2 nM kinesin-1 (F) at 45 minutes. The bar plots were obtained with a binning size of 300 nm. G-I) Simulated kymographs for *C*_*m*_ = 0 (G), 1 (H), and 2 nM (F) showing D (blue), T (red), Tx (green) sites and motor particles (yellow). Each kymograph is accompanied by a zoom on one of its region delimited by the black rectangles. The size of a point is larger than the size of a subunit, so that single sites appear larger than they are. J-L) Simulation lengths distributions for *C*_*m*_ = 0 (J), 1 (K), and 2 nM (L) at 45 minutes. Insets: log-scale plots of the distribution.

Upon addition of kinesin-1 to our assay, the rescue frequency increased with the motor concentration, while the growth and shrinkage velocities as well as the catastrophe frequency of individual microtubules remained unaffected, Fig. 2B,C,E in Ref. (35). Simultaneously, we observed the average microtubule length after 45 min to increase with the kinesin-1 concentration, 13.5 ± 8.1 µm for 1 nM and 19.0 ± 11.6 µm for 2 nM, Figs. 2A-F, S1B-D. For 2 nM kinesin-1 and higher motor concentrations, we did not observe saturation of the average microtubule length within the experimental timeframe of 45 min, Fig. S1C. Note that the standard deviation of the microtubule length distribution at 45 min also increased with the kinesin-1 concentration, Fig. S1D. We hypothesize, that microtubule growth transitions form bounded to unbounded growth (37) at a critical concentration of ≈ 2 nM kinesin-1. An experimental study of the putative transition would require observation times much longer than 45 min.

### The model for rescue at exchange sites reproduces *in vitro* data

In order to theoretically study microtubule length dynamics in presence of molecular motors, we employed stochastic simulations of the model with **R**escue at **E**xchange **S**ites (RES-model), Fig. 1, Methods. Most of the parameter values used in our simulations were determined from our *in vitro* system, Table 1. All simulations started with a single site from which lattices grew, Fig. 2G-I, S2, and S3. Along the lattice, sites in state D could spontaneously change into state Tx and motor particles could further induce such changes, which represent tubulin exchange along a microtubule shaft. A shrinking lattice rescued at Tx sites with a probability *p*_×_. For each condition, we performed *N* = 10^4^ simulations. To compare our simulation results to our experiments, we first determined the lattice length after 45 min simulated time, Figs. 2J-L.

Kymographs obtained from our simulations resembled those from our experiments, Figs. 2G-I, S2, S3. Note that different kymographs for the same conditions varied substantially, Fig. S3. At motor concentration *C*_*m*_ = 0 nM, we observed few rescue events, which occurred at a rate *k*_res_ = 0.5 *µ*m/min, Figs. 2G. This is the same rate as in the experiments. Since growth and shrinkage velocities as well as catastrophe and rescue rates are the same in our experiments and simulations, we expected to obtain the same average lengths. In contrast to our expectation, the experimental average length was larger than the value we obtained in the simulations, Fig. S1A,D. This difference is partly due to the fact that it is difficult to detect microtubules below ∼ 1 µm in our experiments, Fig. 2D, which leads to an overestimation of the experimental average length. However, this does not fully explain the differences between the experimental and theoretical distributions for microtubule lengths lower than 5µm. Additional factors could include molecular processes that we do not account for in our model.

With increasing values of *C*_*m*_, in our simulations, the number of observed rescue events increased, which was also the case in our experiments up to a motor concentration of 2 nM, Figs. 2H,I, S3C-F. Consequently, also the average length increased, Figs. S1B-D. As before, the experimental values were larger than those obtained from the simulations. Concluding, our stochastic simulations reproduce salient features of microtubule growth and length distribution in absence and in presence of motors.

### Average lattice length at steady state

With our simulations we can study the lattice length distribution at steady state. To ensure to reach steady state, we simulated up to 3000 hr, depending on the condition. In line with our results from 45 min simulated time, in steady state, the standard deviation was equal to the average length, Fig. S4. The average lattice length at steady state and the time needed to reach this value increased monotonically with the motor concentration *C*_*m*_, Fig. 3A-C. As the growth and shrinkage velocities and the catastrophe rate were not changed, this increase of the average length was due to a decrease in the fraction of shrinkage periods, Fig. 3D and S5. For *C*_*m*_ ≳ 2.1 nM, the simulations did not reach a steady state.

**Figure 3:**
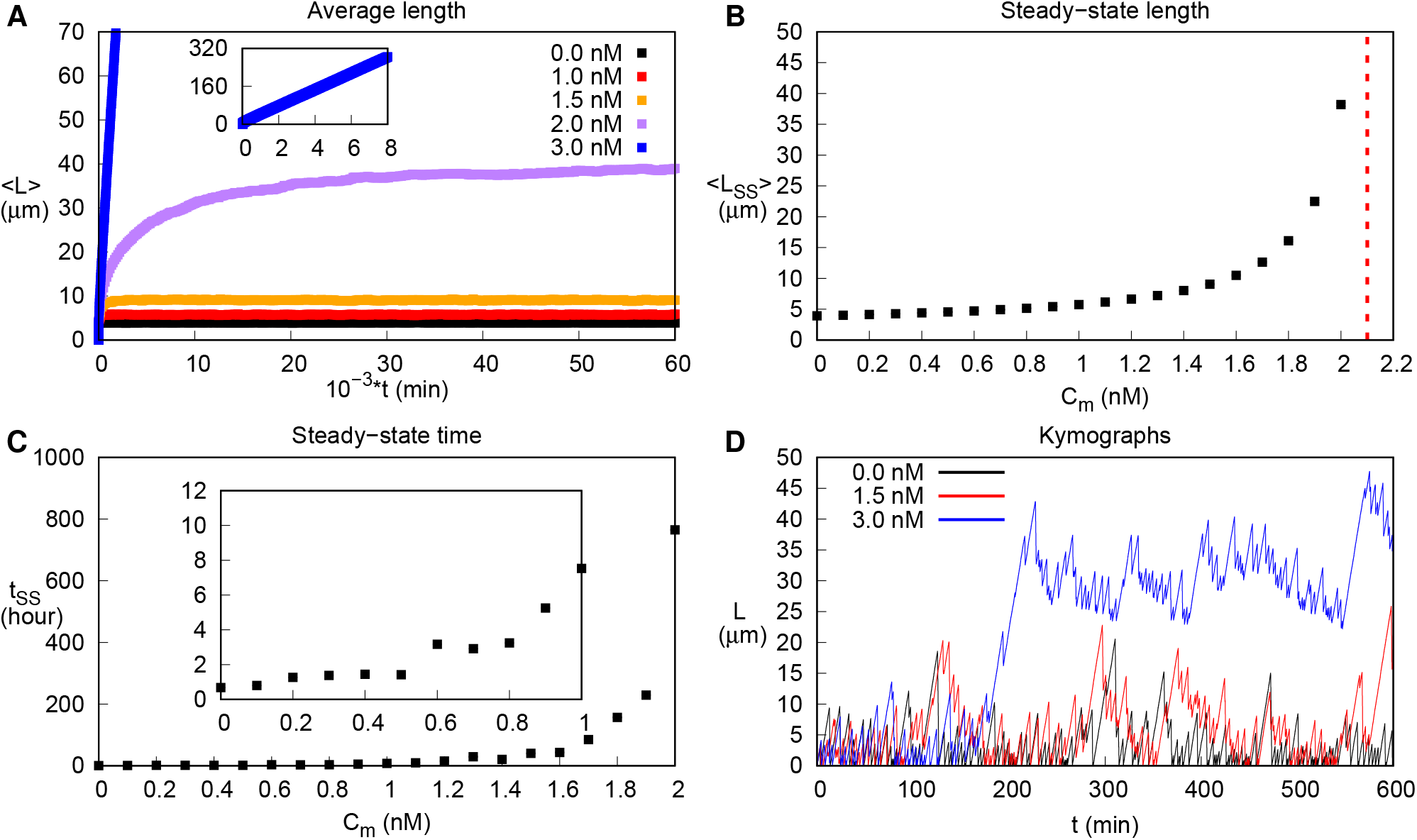
Simulated steady-state average lattice length exhibits a transition. A) Average lattice length as a function of simulation time obtained from 10^4^ repetitions. Inset: complete curve for *C*_*m*_ = 3 nM. B) Average length in steady state as a function of motor concentration. For *C*_*m*_ > 2.1 nM (red dashed line) no steady state was obtained. C) Simulation time needed to reach steady state. The time was recorded when the average length was within one standard deviation from its steady-state value. Inset: zoom for *C*_*m*_ ≤ 1 nM. (D) Long-term length *vs* time for *C*_*m*_ = 0, 1.5 and 3 nM.

### Unbounded lattice growth

In our simulations, for *C*_*m*_ ≥ 2.1 nM, the probability of complete depolymerization *P*_cd_ vanished, because all shrinking lattices rescued, Fig. 4A. As a result, the average lattice length increased at constant velocity and without a bound, Fig. 3A, which was compatible with our experimental data at 2 nM kinesin-1. Indeed, at this kinesin-1 concentration, we could not detect a decrease in the growth velocity of the average microtubule length towards the end of our experiments, Fig. S1C. Together, our experimental and theoretical results indicate that microtubule dynamics exhibits a transition from bounded to unbounded growth at a critical motor concentration of *C*_*m*_ ≈ 2.1 nM. Obviously, in any experiment, microtubule growth would eventually come to a halt, when the tubulin concentration is below the critical concentration of polymerization.

**Figure 4:**
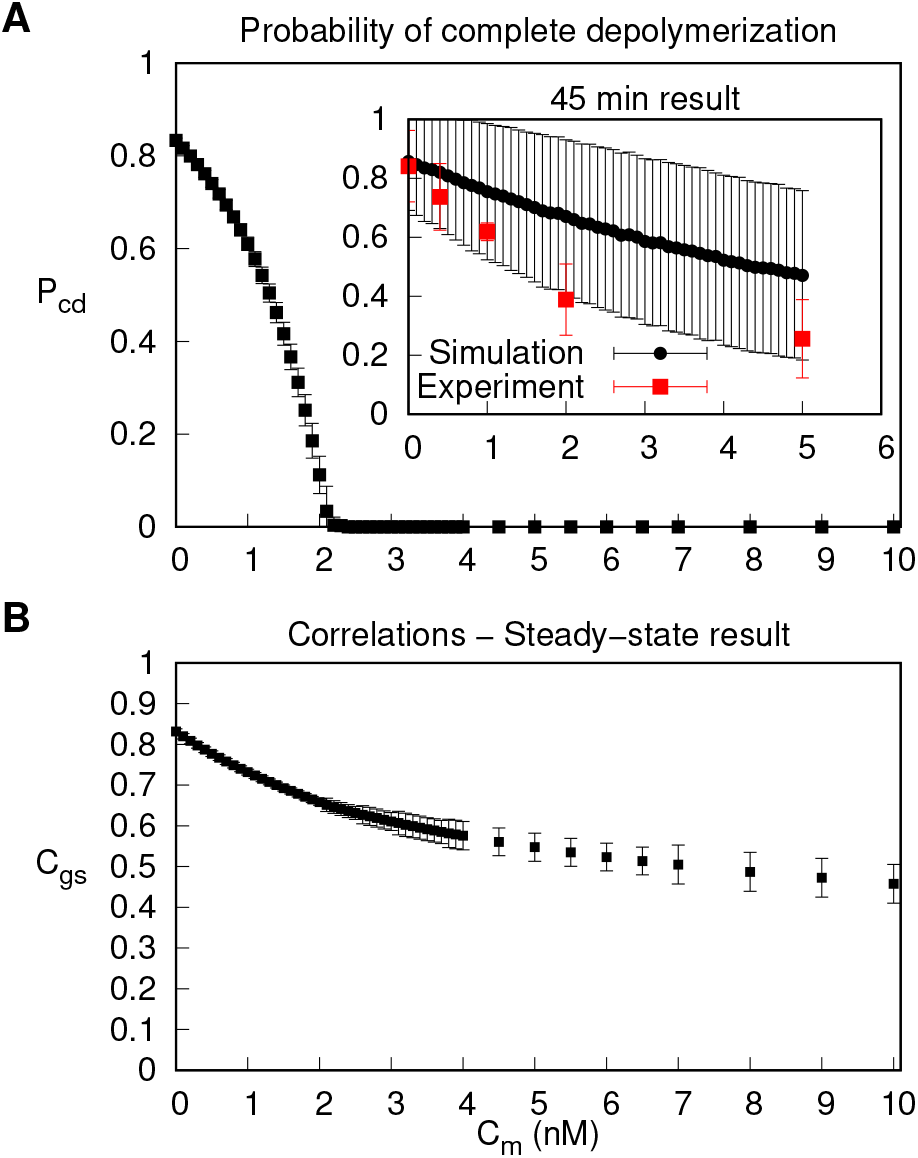
Transition from bounded to unbounded growth regime. A) Probability of complete depolymerization *P*_cd_ as a function of *C*_*m*_. Data points for the bounded growth regime were measured in steady state. Inset: Simulation (black) and experimental data points (red) measured after 45 min. B) Correlation *C*_gs_ between the length of a growth phase and of the subsequent shrinkage phase as a function of *C*_*m*_. In the unbounded growth regime, error bars are larger due to fewer data.

To study the transition from bounded to unbounded growth, we defined the correlation *C*_gs_ between the length added during a growth phase and the length removed during the subsequent shrinkage phase. Explicitly, if *g*_*i*_ denotes the length added during the *i*th growth phase and *s*_*i*_ the length removed during the following shrinkage phase, then the correlation *C*_gs_ is given by

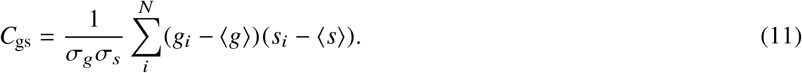

Here, ⟨*g*⟩ and ⟨*s*⟩ denote the respective average lengths added to or removed from the lattice. Furthermore, 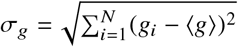 is the standard deviation of the length in the growth phases and, analogously, σ_*s*_ for the shrinkage phases. Finally, *N* is the number of observed pairs of growth and shrinkage phases. The correlation function decreases monotonically with increasing motor concentration *C*_*m*_, Fig. 4B. The correlation function did not exhibit a clear sign of the transition at the critical motor concentration. Below, we come back to this surprising finding.

### Distribution of exchange sites

In our model, rescue can only occur at exchange sites (Tx-state). We thus studied the distribution of the nucleotide states D, T and Tx along the lattice at steady state. For *C*_*m*_ = 0, the probability distribution of T-states, *t*, increased exponentially towards the growing end, whereas the probability distribution of D-states, *d*, showed the opposite behavior, Fig. 5A. The characteristic length of these exponential distributions is essentially given by *v*_*g*_/*k*_hyd_. The probability to find a site in the Tx-state, *t*_×_, was smaller than one percent and decreased linearly over a region that extended from close to the lattice seed to the tip. The non-exponential character of the Tx distribution is a consequence of the spontaneous exchange of D for Tx, the hydrolysis of Tx, and lattice depolymerization.

**Figure 5:**
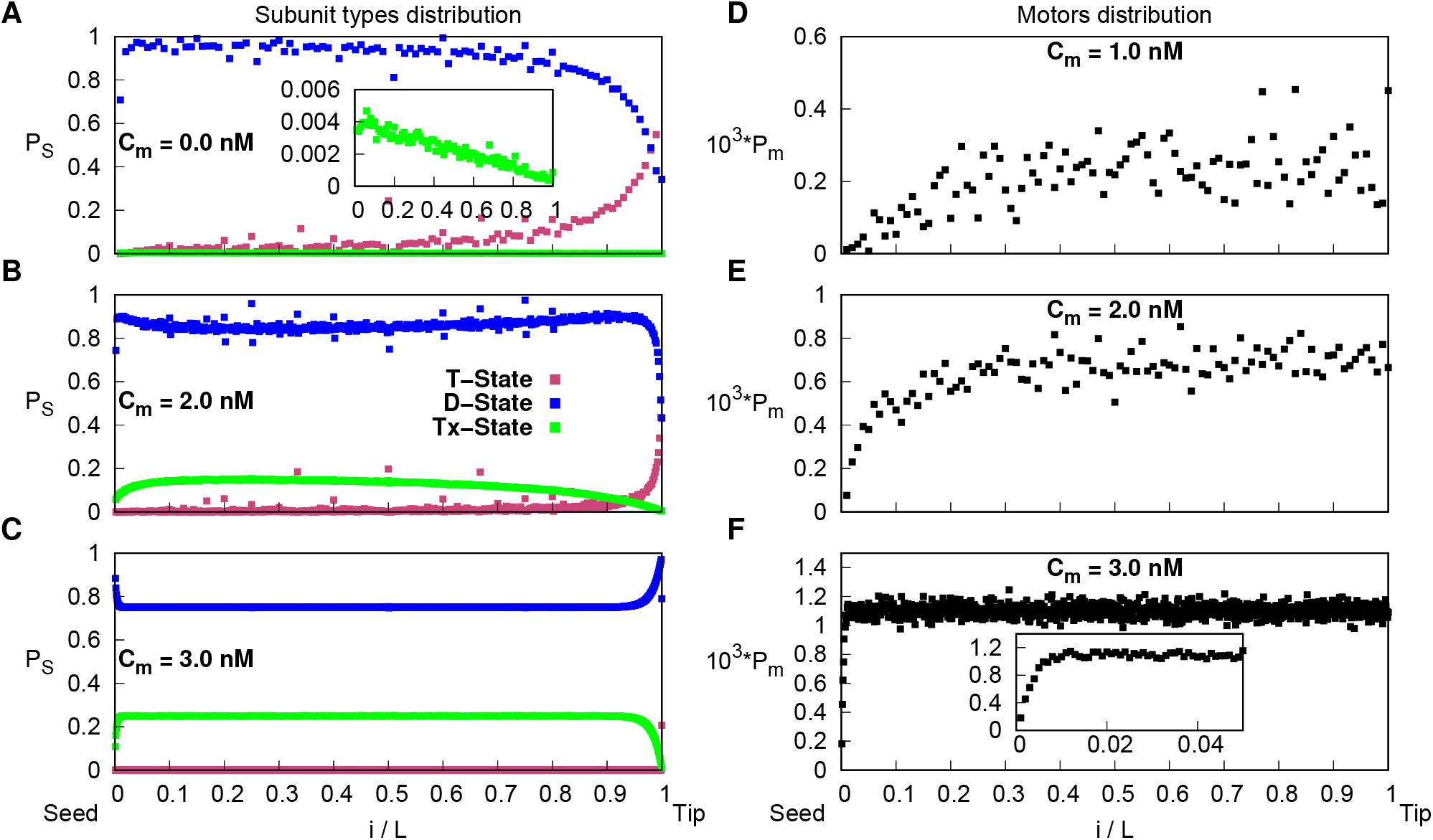
Distribution of states and motors along the lattices. A-C) Average probability distribution of T-(magenta), D-(blue), and Tx-states (green) along the lattice for *C*_*m*_ = 0 (A), 2 (B), and 3 nM (C). Inset: zoom on the Tx-state probability distribution. D-F) Average probability distribution of motors along the lattice *C*_*m*_ = 1 (D), 2 (E) and 3 nM (F). Inset: zoom on the distribution close to the seed. The binning size for the distributions is 10^−2^ (A,D,E) and 10^−3^ (B,C,F).

In the presence of motors, the distribution *t* was not altered, Fig. 5B,C. In contrast, the steady-state distribution *d* was reduced in the bulk of the lattice compared to the boundaries, Fig. 5B. Since motors were homogenously distributed sufficiently far away from the lattice seed, Fig. 5D,E, one can attribute this decrease of *d* to an effective increase of the exchange rate. Accordingly, the decrease in *d* was compensated by an increase of *t*_×_. We note that although the probability to find a motor on a given site is very small, < 0.001, the probability *t*_×_ increases by a factor of ∼ 40 for *C*_*m*_ = 2 nM compared to *C*_*m*_ = 0 nM. In the unbounded growth regime and relative to the lattice length, the regions of the exponential decay of *t* and the corresponding increase of *d* asymptotically shrank to zero and we neglect it in the following discussion. The distribution *d* decreased from the growing end towards a plateau value that is given by

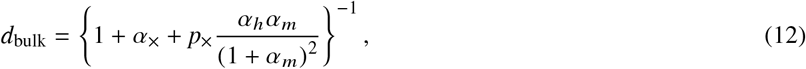

where *α*_×_ = *k*_×_ /*k*_hydX_, *α*_*h*_ = *k*_hop_/*k*_hydX_, and *α*_*m*_ = *k*_mot,d_/*k*_mot,a_, see below. Since on most sites *t* ≈ 0, we have *t*_×_ = 1 − *d*.

In summary, in the absence of motors, the probability that a site is in the Tx-state increases with the time the site has spent in the lattice, Fig. 5A. Since shrinking lattices can rescue at exchange sites, the probability to rescue is thus position-dependent. With increasing motor concentration, this probability approaches a constant value along the lattice and rescues occur at a constant rate.

### Effective rescue rate

As a consequence of the position-dependent rescue rate, the distribution of lengths removed between a catastrophe and a rescue event was not exponential, Fig. 6A. Let us recall that we do not consider regrowth from the seed to be a rescue event. With increasing motor concentration *C*_*m*_, we observed a decrease of the typical rescue length relative to the lattice length, Fig. 6A.

**Figure 6:**
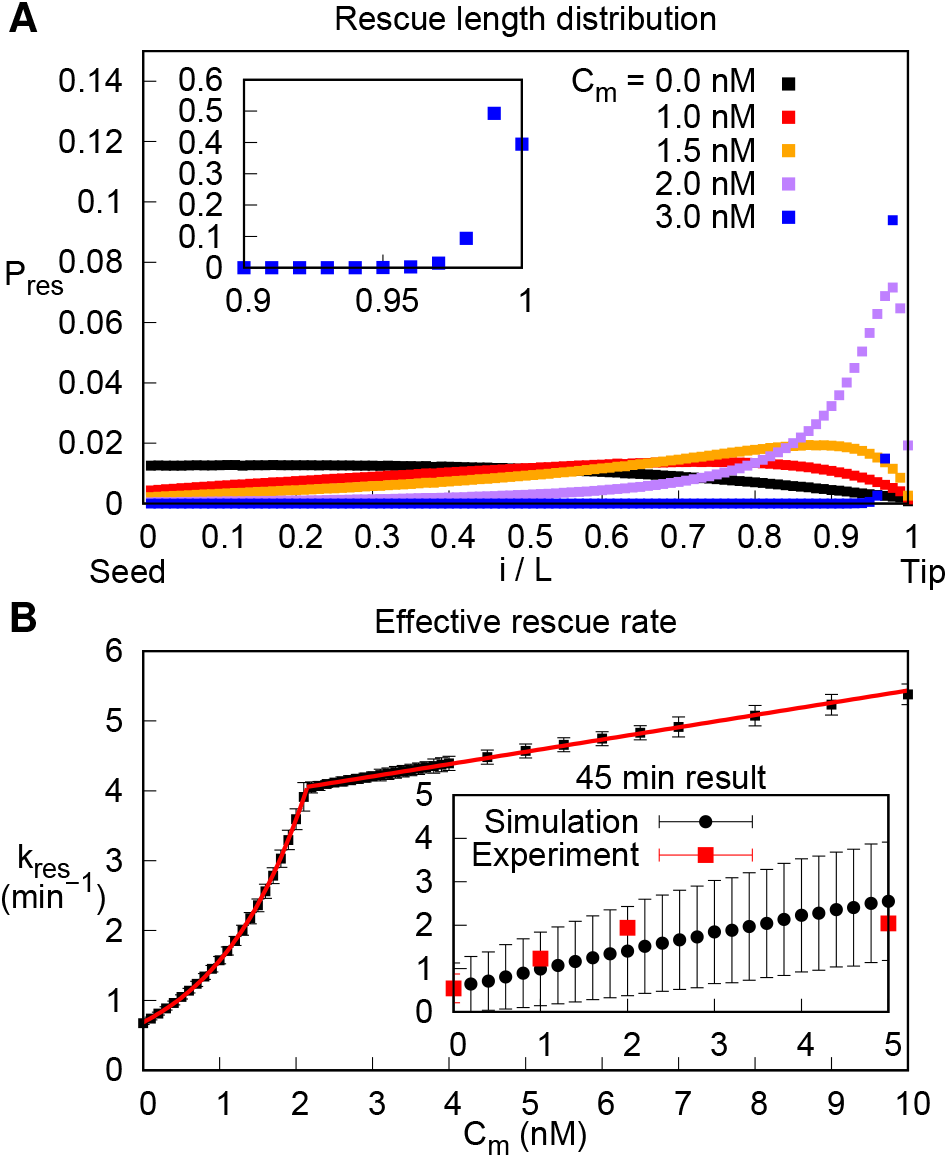
Effect of motors on rescues. A) Probability distributions of the shrinkage length relative to the lattice length prior to depolymerization for different motor concentrations. For 3 nM the peak shifts towards the tip as the simulation time increases. Inset: zoom for 3 nM. B) The effective rescue rate *k*_res_ as a function of *C*_*m*_. Red line: exponential fit up to *C*_*m*_ = 2.1 nM and a linear fit for *C*_*m*_ > 2.1 nM. Inset: experimental data and simulation results at 45 min.

Within our *in vitro* experiments it was not possible to measure the position-dependent rescue rate. Instead, we measured an effective position-independent rescue rate to quantitatively characterize the microtubule dynamics in presence of kinesin-1 (35). To this end, the total number of rescue events was divided by the total time spent in the shrinking state for all microtubules observed under one condition. In our experiments, this effective rescue rate increased with the kinesin-1 concentration, Fig. 6B, inset.

We obtained the effective rescue rate also from our simulation data for all motor concentrations. For *C*_*m*_ ≤ 1 nM, the effective rates from our simulations are close to the experimentally measured values. For larger values of *C*_*m*_, the experimental effective rescue rates are smaller than those from the simulations at steady state. Since we know that in our simulations the steady state is not reached after 45 min simulated time for *C*_*m*_ > 1 nM, we also measured the effective rescue rate from simulations of 45 min simulated time. In that case, we obtained a good match between our experimental and our simulation rates, Fig. 6B, inset. At steady state the effective rescue rate increased with *C*_*m*_, Fig. 6B. As a function of the motor concentration *C*_*m*_ the effective rescue rate increased exponentially from *C*_*m*_ = 0 to the critical motor concentration, *k*_res_ ∝ exp *C*_*m*_/*C*_*m,c*_ with *C*_*m,c*_ = 1.209 ± 0.004 nM. Beyond this concentration the function increased linearly with a slope *s* = 0.175 ± 0.001 min^−1^nM^−1^. The transition from exponential to linear increase occurred at the same value of *C*_*m*_ as the transition from bounded to unbounded lattice growth, Fig. 4A.

The transition between bounded and unbounded growth can also be induced by increasing the spontaneous exchange rate *k*_×_, Fig. S6. Independently of whether the transition was induced by increasing the motor concentration or the spontaneous exchange rate, it exhibited the same features. The only notable difference is the distribution of Tx close to the seed, Fig. S7, which has an impact on the average lattice length in steady state.

### The average microtubule dynamics is captured by the effective rescue rate

Having introduced the effective rescue rate, we wondered how the lattice dynamics changes if rescue events occur at a constant effective rate *k*_res_, instead of being limited to exchanged sites. This dynamics was studied by Dogterom and Leibler (37) and we will refer to it as the DL-model. In addition to *k*_res_, the DL-model depends only on the growth velocity *v*_*g*_ ≡ *ak*_*a*_, the shrinkage velocity *v*_*s*_ ≡ *ak*_*d*_, and the catastrophe rate *k*_cat_.

As long as *v*_*g*_ *k*_res_ < *v*_*s*_ *k*_cat_, the system reaches a steady state with an average length ⟨*L*⟩_DL_, which is given explicitly by (37)

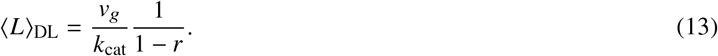

Here *r* = *v*_*g*_ *k*_res_/(*v*_*s*_ *k*_cat_), which is the only independent control parameter in the DL-model. In the unbounded growth regime, the average growth velocity ⟨*v*_g_⟩_DL_ is given by (37)

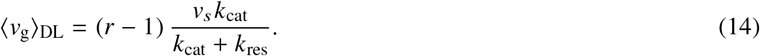

In the DL-model, the critical value of the control parameter is *r*^∗^ = 1. For the values of the parameters *k*_*a*_, *k*_*d*_, and *k*_cat_ used in our simulations, we obtain 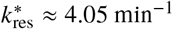. Remarkably, this value matches the effective rescue rate at the transition from bounded to unbounded growth in our model – independently of whether the transition was induced by changing the motor concentration *C*_*m*_ or the spontaneous exchange rate *k*_×_, Fig. 6B, S6D.

Above, we have characterized the transition to unbounded growth by the probability for complete depolymerization, *P*_cd_. In the following, we compute this value for the DL-model. In the bounded growth regime, this probability can be written as

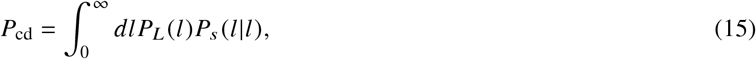

where *P*_*L*_ is probability distribution of the lattice length and *P*_*s*_ (*s* | *l*) the probability distribution that a lattice shrinks by a length *s*, when it had a length *l* at the catastrophe event. In the DL-model, we have

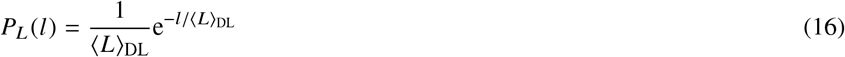

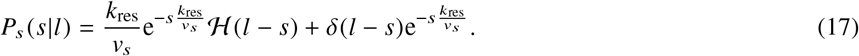

Here, *δ* denotes Dirac’s *δ*-distribution with *δ*(*x*) = 0 for *x ≠* 0 and ∫ *dx δ*(*x*) = 1, and ℋ denotes the Heaviside step function with ℋ (*x*) = 1 for *x* > 0 and ℋ (*x*) = 0 for *x* ≤ 0. Calculating the integrals in Eq. (15) yields

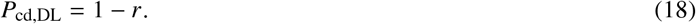

Note that this expression was obtained for the steady state and thus only valid for *r* ≤ 1.

In the unbounded growth regime, the lattice grows indefinitely. Therefore, *P*_*L*_ (*l, t*) tends to zero for finite *l* and *t* → ∞. Consequently, *P*_cd_ = 0. Our analytical result agrees with simulations of the DL-model, Fig. 7A. For the RES-model, which accounts for spontaneous and motor-induced exchange, Eq. (18) also describes *P*_cd_ if we use the effective rate *k*_res_ to calculate *r*, Figs. 6B, 7A, S6D. In the unbounded growth regime, the average growth velocity *v*_*g*_, Eq. (14), yields the corresponding value for the RES-model, if *r* is again obtained from the effective rate *k*_res_, Fig. 7B.

**Figure 7:**
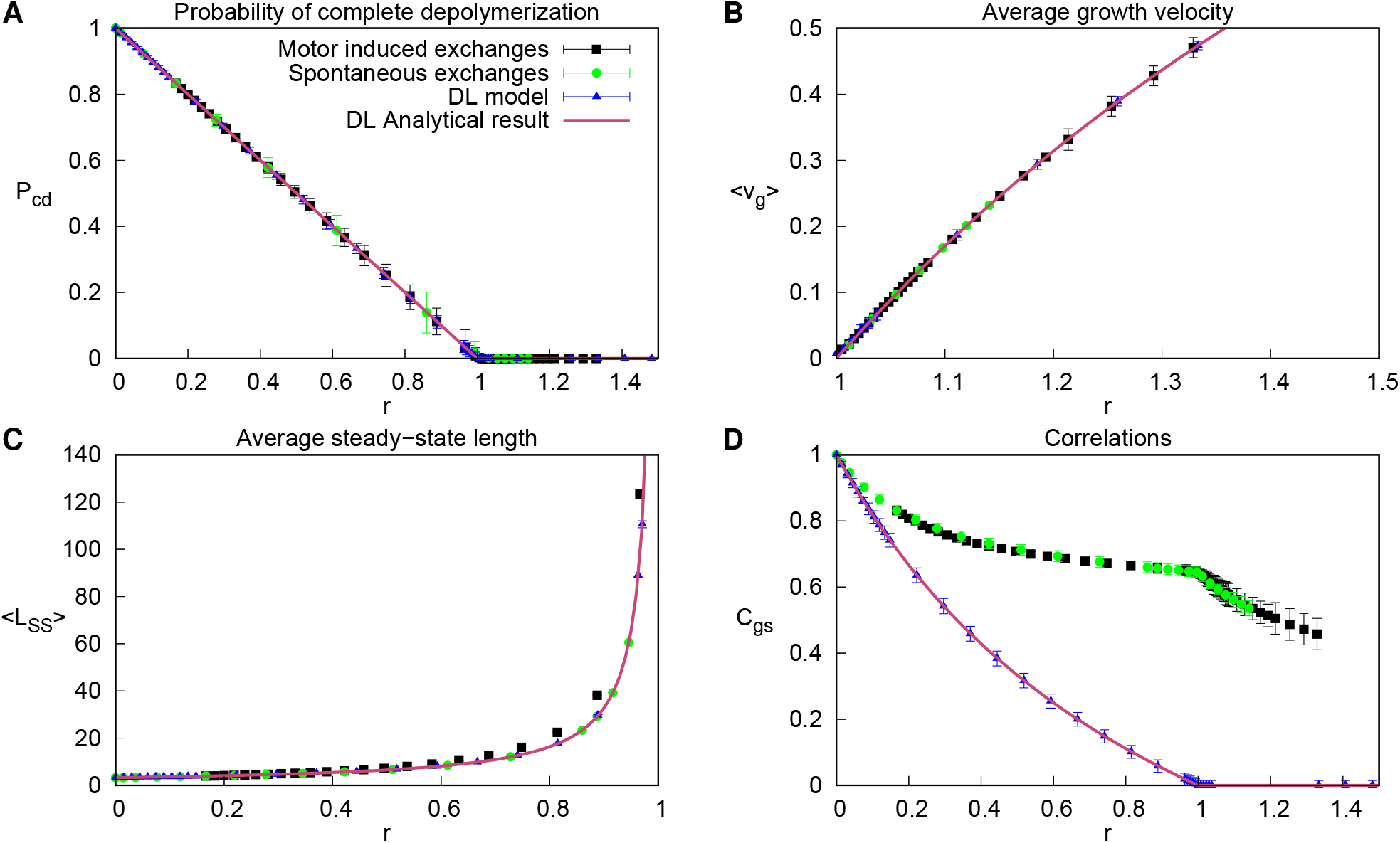
Comparison of our RES-model to the DL-model. A) Probability of complete depolymerization *P*_cd_ as a function of *r* = *k*_a_*k*_res_/*k*_d_*k*_cat_ with *k*_res_ from Fig. 6B. B) The average growth velocity in the unbounded growth regime as a function of *r*. C) Average length in the steady state of the bounded growth regime as a function of *r*. D) The correlation *C*_gs_ as a function of *r*.

In the bounded growth regime, after computing *r* as before, Eq. (13) gives the average lattice length for the RES-model accounting only for spontaneous exchange, Fig. 7C. If also motor-induced exchanges are considered, we find that Eq. (13) underestimates the average length obtained from the simulations, Fig. 7C. This difference can be explained as follows: adjacent to the seed, the probability to find Tx sites is significantly smaller than in case of a constant (effective) spontaneous exchange rate, Figs. 5B and S7B. Consequently, the lattice needs to be longer to achieve the same effective rescue rate. We conclude that the average behavior of the RES-model is captured by the DL-model if the control parameter *r* is computed using the effective rescue rate *k*_res_, Figs. 6B and S6D.

In striking contrast, as a function of *r*, the correlation *C*_gs_ differs strongly between the models, Fig. 7D. Whereas the correlation function for the DL-model drops to zero at the transition, it stays finite for the RES-model. Still, as a function of *r, C*_gs_ displays a clear sign of the transition, which is not the case if considered as a function of *C*_*m*_ or *k*_×_, Figs. 4B and S6B. Also, the functional dependence of *C*_gs_ on *r* is the same for the cases with and without motors.

We can calculate the correlation function for the DL-model in the bounded growth regime. To this end we write the correlation function as

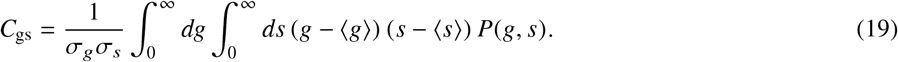

Here, ⟨*g*⟩ = *v*_*g*_/*k*_cat_ and ⟨*s*⟩ = *v*_*g*_/*k*_cat_ are the expectation values of the growth and shrinkage lengths, σ_*g*_ = ⟨*g*⟩ and σ_*s*_ = ⟨*s*⟩ the corresponding standard deviations, and *P*(*g, s*) is the probability distribution for observing growth by a length *g* and a subsequent shrinkage by a length *s*. It can be written as

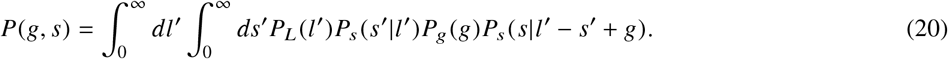

Here, *P*_*g*_ is the probability distribution for the length of a growth phase. Explicitly,

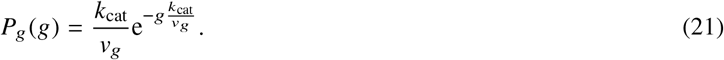

Injecting Eqs. (16), (17), (20) and (21) into Eq. (19) yields

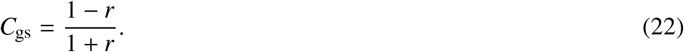

In the limiting cases *r* = 0 and *r* = 1, we have *C*_gs_(*r* = 0) = 1 and *C*_gs_(*r* = 1) = 0 as expected. In the unbounded growth regime, *r* ≥ 1, the growth and shrinkage phases are independent of each other and *C*_gs_ = 0. The simulation results match Eq. (22), Fig. 7D.

### Meanfield analysis

We study the lattice dynamics using a meanfield approach by expressing all correlation functions in terms of products of mean values, Eqs. (1)-(5), Methods. We obtained the steady-state solutions of these equations by using an explicit Euler scheme for the time evolution, Fig. 8. In absence of motor particles and for sufficiently small values of the spontaneous exchange rate *k*_×_, the meanfield solutions agreed well with our simulation results, Figs. 8A-D, S8A-D. Only the distribution of Tx-states differed significantly between these two approaches, Figs. 8C, S8C. This difference increased with increasing *k*_×_, Fig. S8E-H, and was due to the tight link between rescue events and the presence of sites in the Tx-state.

**Figure 8:**
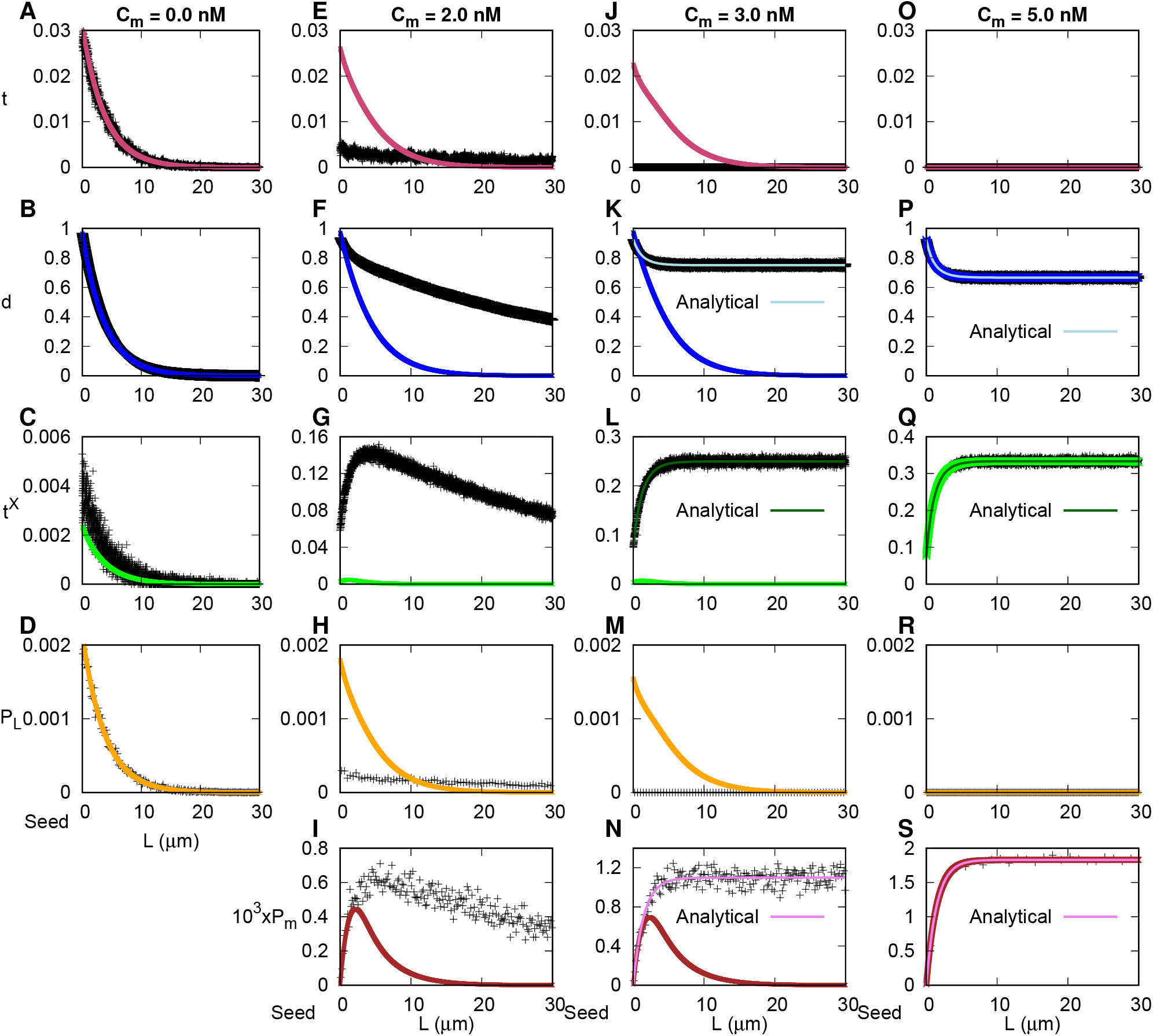
Meanfield solutions for different motor concentrations *C*_*m*_. Distributions of T-(A, E, J,O), D-(B, F, K, P), Tx-states (C, G, L, Q), lattice lengths (D, H, M, R), and motors (I, N, S) with respect to the lattice seed for *C*_*m*_ = 0 (A-D), 2 (E-I), 3 (J-N), and 5 nM (O-S). Black crosses: simulation results; dark blue, light green, orange, and magenta lines: numerical solutions to the meanfield equations; light blue, dark green, and purple lines: analytical solutions to the meanfield equations.

For *C*_*m*_ ≠ 0, the effects of these correlations were stronger than for *C*_*m*_ = 0, increasing the difference between the simulations and the meanfield equations, Fig. 8E-H. Similarly, the motor distributions from the simulations and the meanfield equations differed sufficiently far away from the seed, Fig. 8I. For *C*_*m*_ = 3.0 nM, our stochastic simulations differed qualitatively from the meanfield equations as the former was in the unbounded regime, whereas it was still in the bounded regime for the latter,

Fig. 8J-N. Indeed, the critical value of *C*_*m*_ for the transition from bounded to unbounded growth was *C*_*m*_ ≈ 2.1 nM in the stochastic simulations and *C*_*m*_ ≈ 4.5 nM in the meanfield approach. Let us also note that the transition to unbounded growth occurred at different values of the spontaneous exchange rate *k*_×_ in the stochastic simulations and the meanfield equations. In this case, the transition occurred for smaller values in the meanfield approach, Fig. S8I-L. For *C*_*m*_ = 5.0 nM, simulations and meanfield equations yielded unbounded growth and the distributions were in perfect agreement, Fig. 8O-S. The same was true in the unbounded regime, when motor particles were absent, Fig. S8M-O.

In the unbounded growth regime, we could compute the various distributions analytically by assuming that the lattice is infinitely long. In that case, we took the continuum limit, Methods, and solved Eqs. (9)-(10). Explicitly, we found for the steady state:

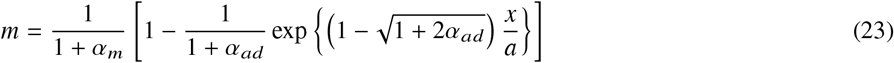

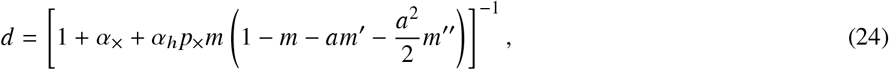

where *α*_*ad*_ = (*k*_mot,a_ + *k*_mot,d_)/*k*_hop_, and *t*_×_ = 1 − *d*, Methods. This solution exactly matched the distribution close to the seed obtained from our simulations, Fig. 8J-S, S8M-O.

Finally, we note that it follows from Eqs. (9) and (24) that the effect of the motors on the distribution can be exactly mapped to a position-dependent exchange rate

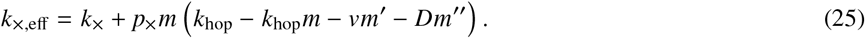

## CONCLUSION

Our analysis shows that molecular motors can significantly change the average length of microtubules by inducing tubulin exchange along the shaft. This observation holds true even if motors are small in number and when only a small fraction of motor steps leads to tubulin exchanges. As a consequence, in cells, microtubules used for transport are stabilized such that transport can lead to microtubule network reorganization. This mechanism offers a direct link between local extracellular stimuli and cell polarization.

Remarkably, although the average effects of molecular motors on microtubule dynamics can be captured by an effective rescue rate, the dynamics of individual microtubules differs dramatically if rescues are governed by an effective rate or occur only at tubulin exchange sites. This is evident when considering the correlation between growth and subsequent shrinkage lengths. Future work will address possible implications for cellular processes and microtubule network organization.

Our simulations focus on the study of microtubule length dynamics in presence of motor-induced tubulin exchange. For this reason it neglects salient aspects of the microtubule tip dynamics, in particular, the link between the ’GTP-cap’, a region of GTP tubulin at a growing microtubule end, and catastrophes. It will be interesting to see, how the detailed tip dynamics affects the correlation between growth and shrinkage lengths. Furthermore, we assume a reservoir of tubulin dimers. Future studies will address the question how a limited pool of tubulin dimers impacts on the behavior of the system.

## AUTHOR CONTRIBUTIONS

KK and CA designed the research. JS carried out all simulations, JS analyzed the data. MAC performed the experiments, JS, KK, and CA wrote the article.

## ACKNOWLEDGMENTS

We thank M. Gonzalez-Gaitan for discussions.

## SUPPLEMENTARY MATERIAL

An online supplement to this article can be found by visiting XXX

**Figure S1:**
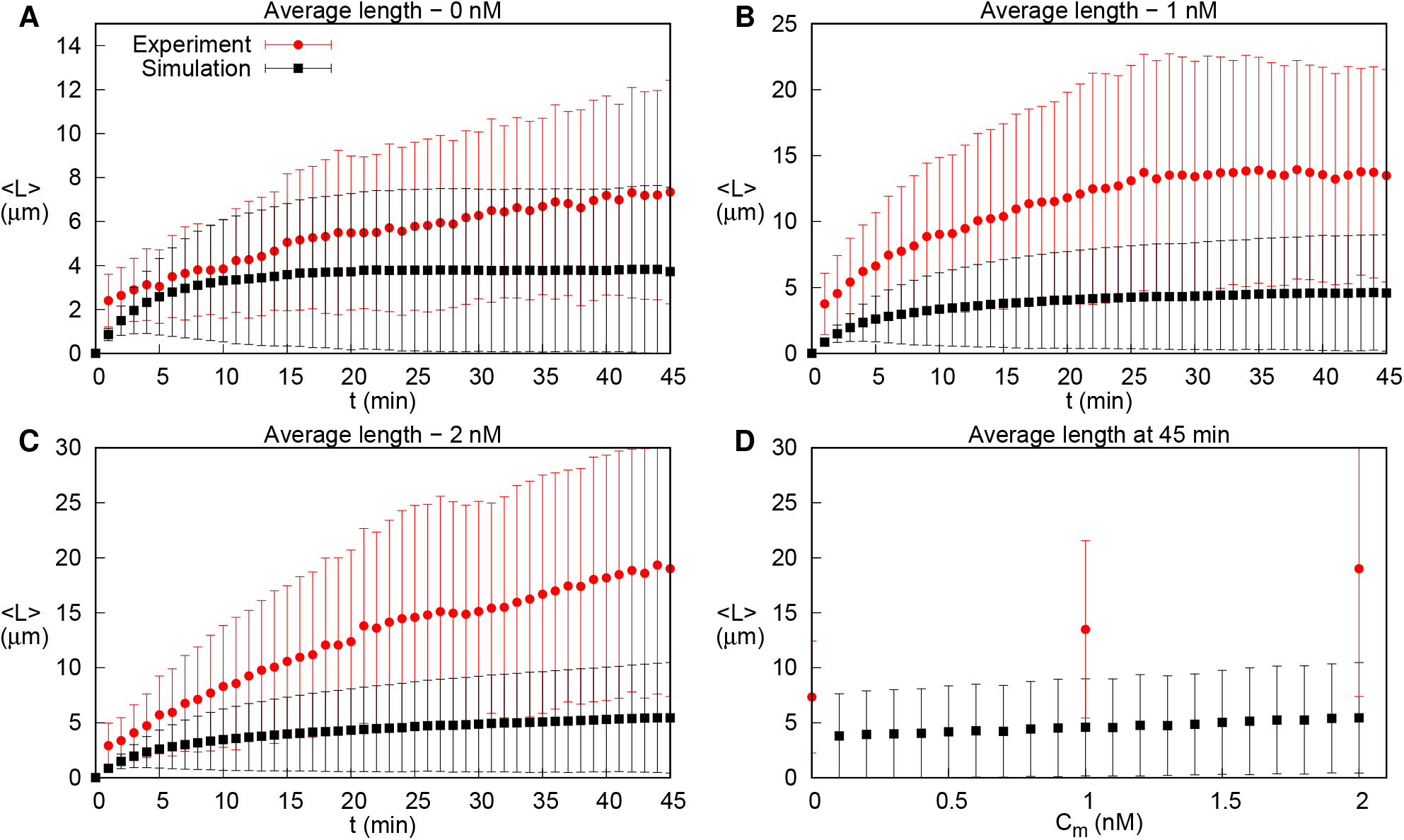
Average length form experiment (red) and simulations (black). A-C) Average length as a function of time for motor concentrations *C*_*m*_ = 0 nM (A), *C*_*m*_ = 1 nM (B), and *C*_*m*_ = 2 nM (C). D) Average microtubule length at 45 min as a function of *C*_*m*_.

**Figure S2:**
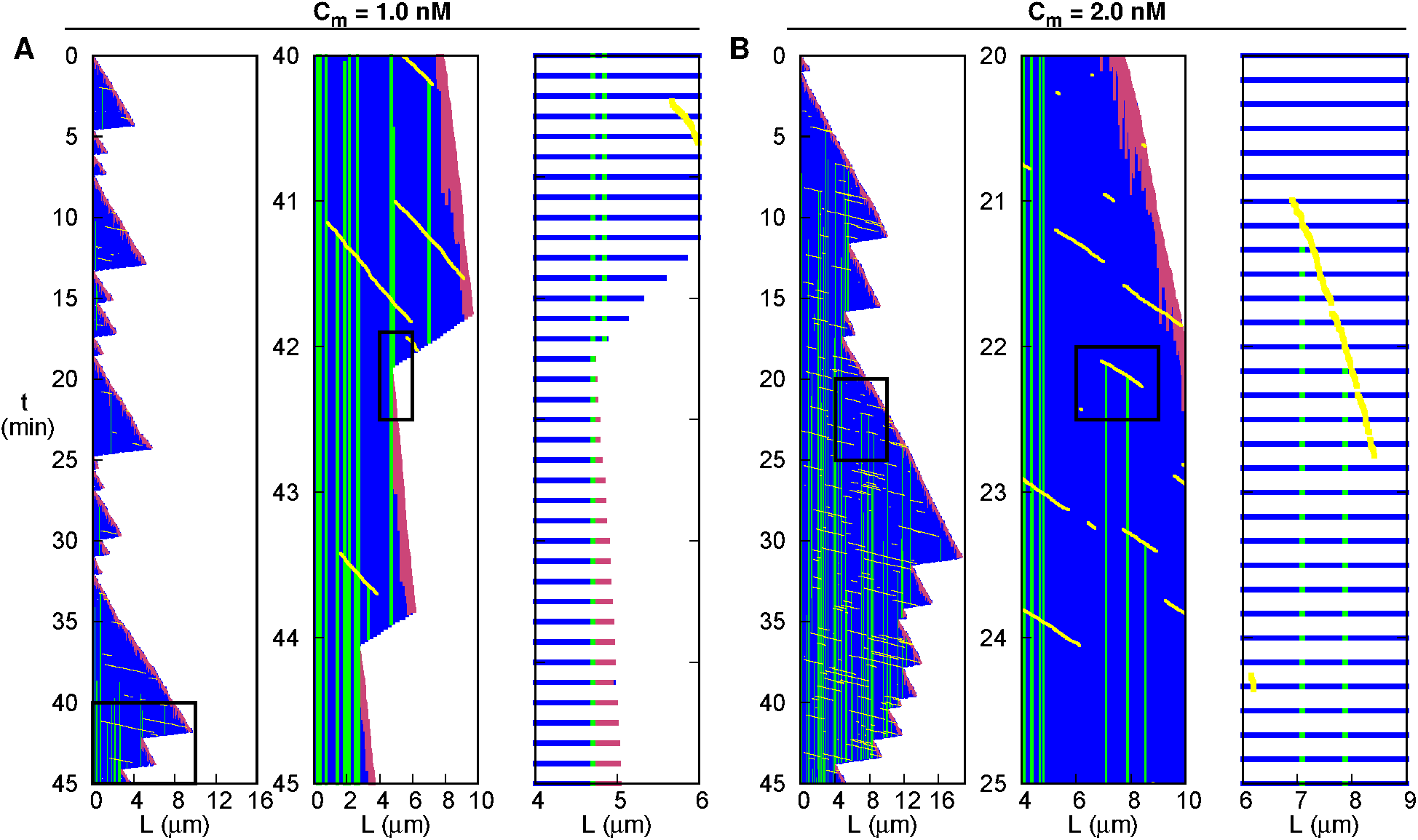
Site state dependent rescue and motor induced exchange. Kymographs of simulated microtubules for 45 min with *C*_*m*_ = 1 (A) and 2 nM (B) showing D (blue), T (magenta), Tx (green) sites and motor particles (yellow). Each kymograph is accompanied by successive zooms (black rectangle) exhibiting a rescue event (A) and motor induced exchange (B).

**Figure S3:**
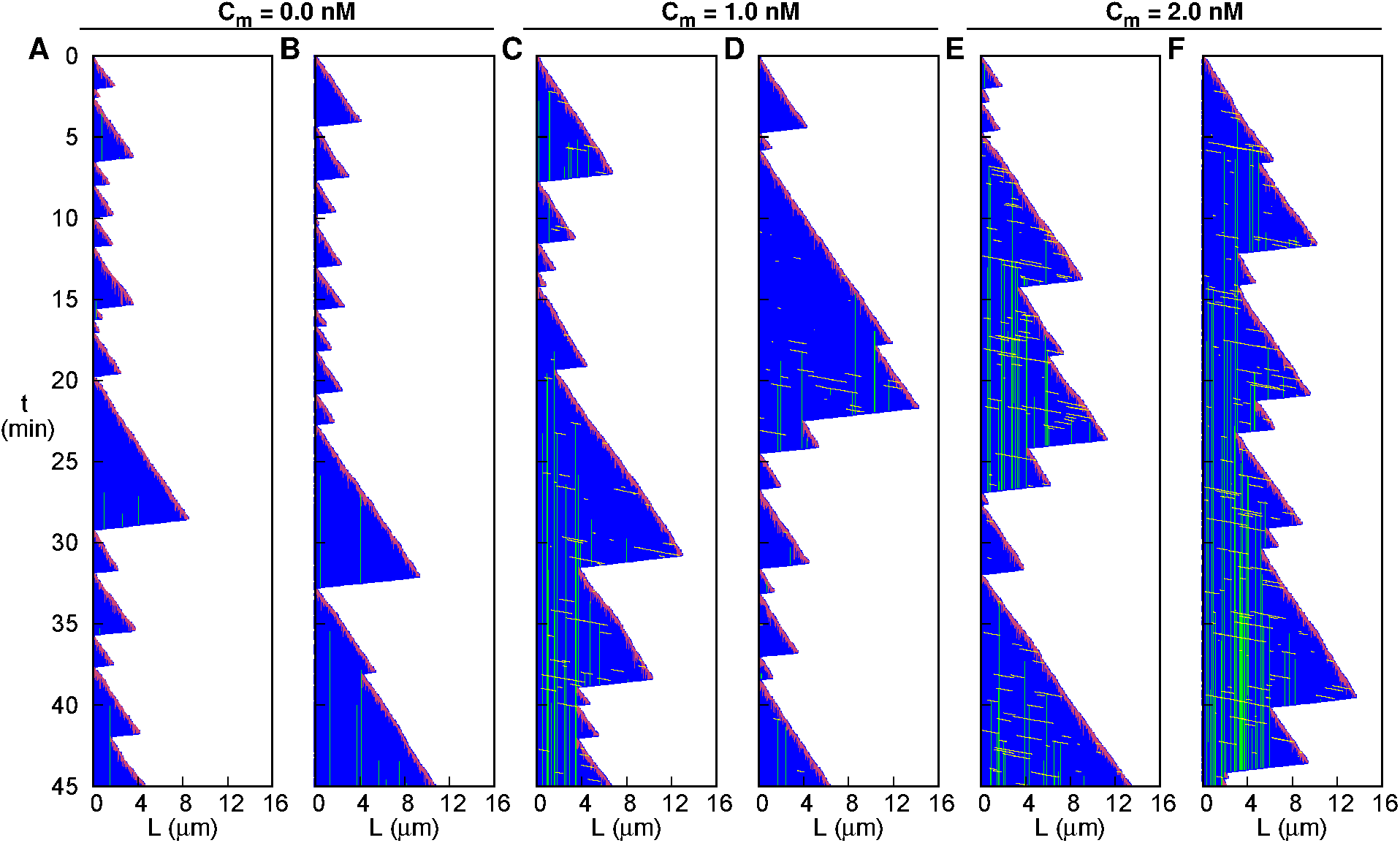
Fluctuations in individual simulated microtubule trajectories during 45 min. Kymographs of simulated microtubules with *C*_*m*_ = 0 (A-B), 1 (C-D), and 2 nM (E-F) showing D (blue), T (magenta), Tx (green) sites and motor particles (yellow).

**Figure S4:**
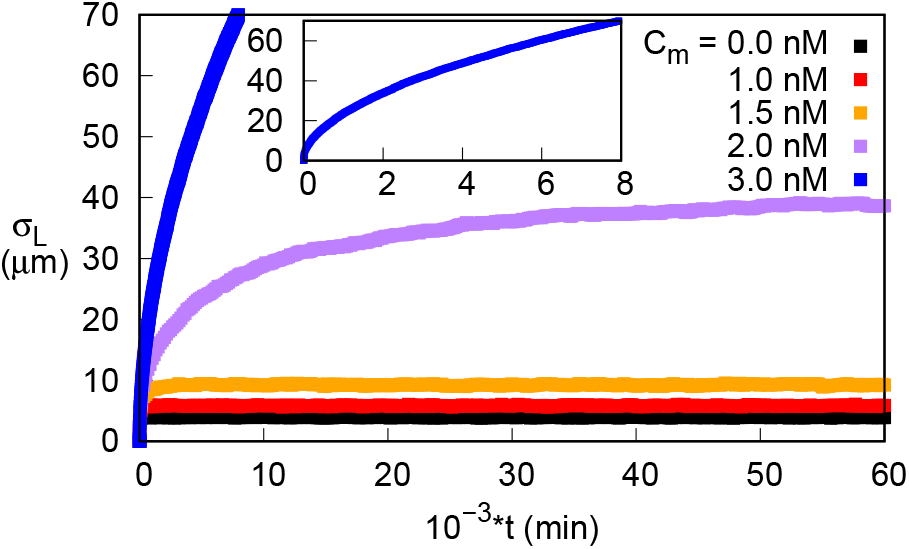
Standard deviation of the length as function of the simulated time for *C*_*m*_ = 0, 1, 1.5, 2 and 3 nM. In the bounded growth regime, *C*_*m*_ = 0, 1, 1.5 and 2 nM, the standard deviation is equal to its average, indicating an exponential distribution. In the unbounded growth regime, *C*_*m*_ = 3 nM, the standard deviation is not equal to the average

**Figure S5:**
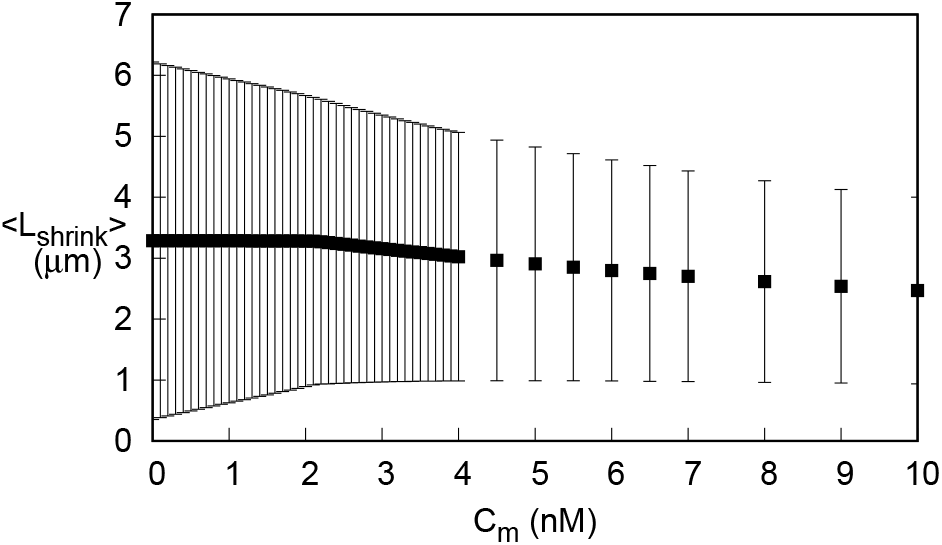
Average length of the shrinkage epsiodes as a function of the motors concentration. Error bars represent standard deviation. In the bounded growth regime, up to *C*_*m*_ = 2.1 nM, the average length of the shrinkage phases stays constant. In the unbounded growth regime, *C*_*m*_ > 2.1 nM, the average shrinkage length decreases with the motor concentration.

**Figure S6:**
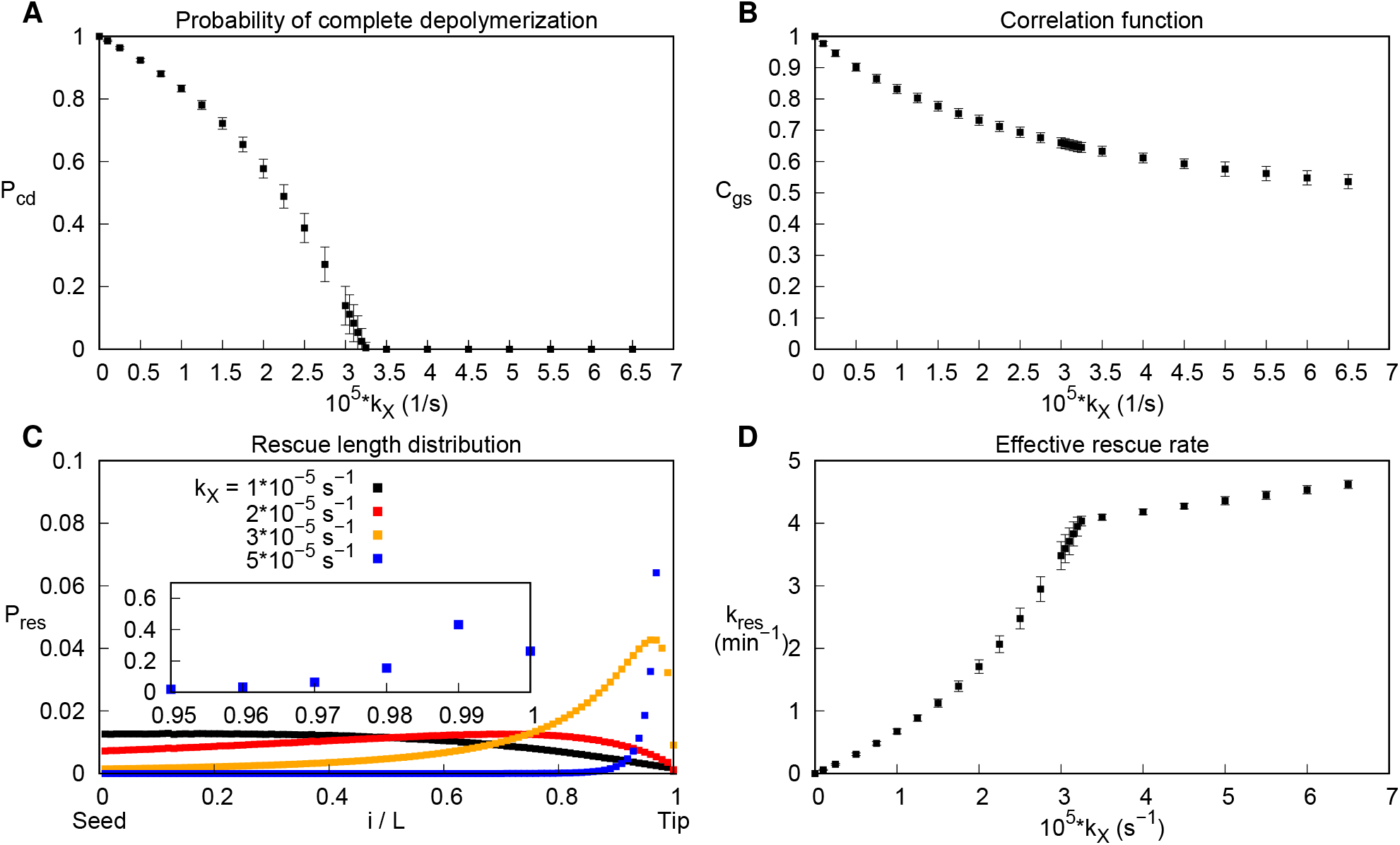
Dependence on the spontaneous exchange rate *k*_×_. A) Probability of complete depolymerization *P*_cd_. Data points for the bounded growth regime were measured in the steady state. B) Correlation *C*_gs_ between the length of a growth phase and of the subsequent shrinkage phase. In the unbounded growth regime error bars are larger due to fewer data. C) Probability distributions of the shrinkage length relative to the microtubule length prior to depolymerization. For 5 × 10^−5^ *s*^−1^ the peak shifts towards the tip as the simulation time increases. Inset: zoom for 5 × 10^−5^ *s*^−1^. D) The effective rescue rate *k*_res_.

**Figure S7:**
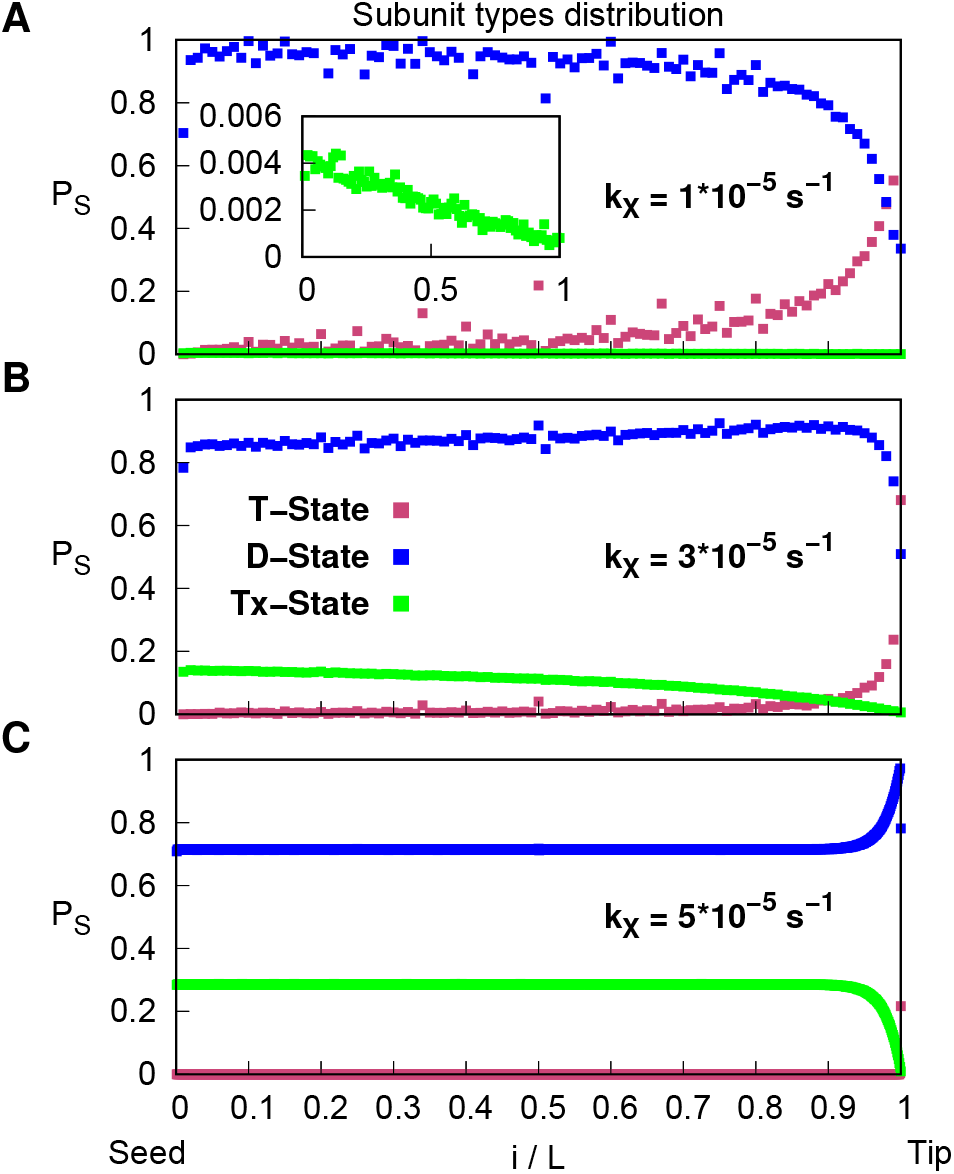
Distribution of subunit types in presence of spontaneous exchanges only. Average probability distribution of T-(magenta), D-(blue), and Tx-subunits (green) along the microtubule for *k*_×_ = 1 × 10^−5^ (A), 3 × 10^−5^ (B), and 5 × 10^−5^ *s*^−1^ (C). The inset in (A) is a zoom on the Tx probability distribution.

**Figure S8:**
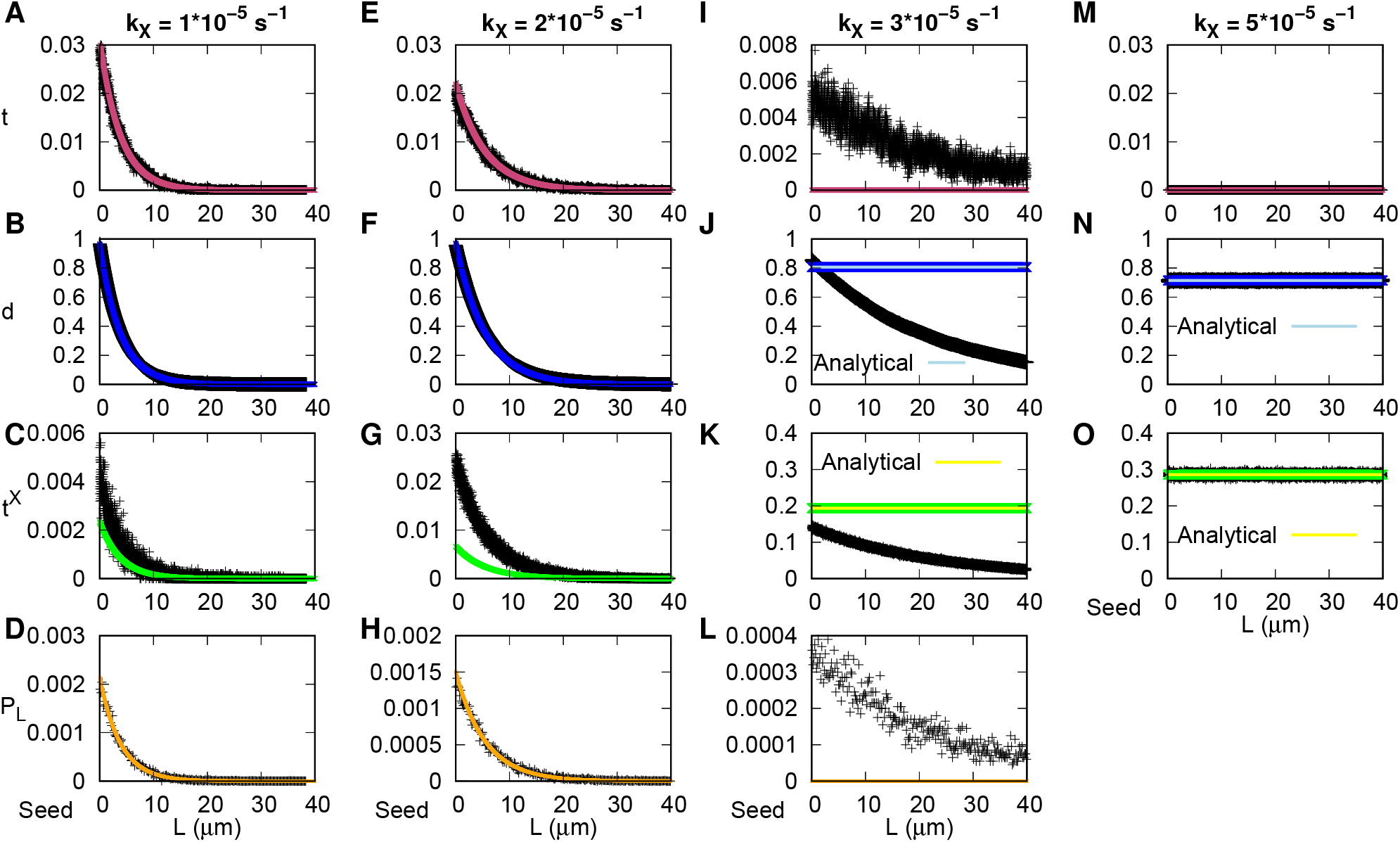
Meanfield solutions for different spontaneous exchange rates *k*_×_. Distributions of T-(A, E, I, M), D-(B, F, J, N), Tx-states (C, G, K,O), lattice lengths (D, H, L), with respect to the lattice seed for *k*_×_ = 1 × 10^−5^ (A-D), 2 × 10^−5^ (E-H), 3 × 10^−5^ (I-L), and 5 × 10^−5^ *s*^−1^ (M-O). Black crosses: simulation results; dark blue, light green, orange, and magenta lines: numerical solutions to the meanfield equations; light blue and yellow lines: analytical solutions to the meanfield equations.

## Notes

### Competing Interest Statement

The authors have declared no competing interest.

